# Prospective role of PAK6 and 14-3-3γ as biomarkers for Parkinson’s disease

**DOI:** 10.1101/2023.04.28.538525

**Authors:** Elena Giusto, Lorenza Maistrello, Lucia Iannotta, Veronica Giusti, Ludovica Iovino, Rina Bandopadhyay, Angelo Antonini, Luigi Bubacco, Rita Barresi, Nicoletta Plotegher, Elisa Greggio, Laura Civiero

## Abstract

2.

**Background:** Parkinson’s disease is a progressive neurodegenerative disorder mainly distinguished by sporadic aetiology, although a genetic component is also well established. Variants in the *LRRK2* gene are associated with both familiar and sporadic disease. We have previously shown that PAK6 and 14-3-3γ protein interact with and regulate the activity of LRRK2.

**Objectives:** The aim of this study is to quantify PAK6 and 14-3-3γ in plasma as a reliable biomarker strategy for the diagnosis of both sporadic and LRRK2-linked Parkinson’s disease.

**Methods:** After an initial quantification of PAK6 and 14-3-3γ expression by means of Western blot in post-mortem human brains, we verified the presence of the two proteins in plasma by using quantitative ELISA tests. We analysed samples obtained from 39 healthy subjects, 40 patients with sporadic Parkinson’s disease, 50 LRRK2-G2019S non-manifesting carriers and 31 patients with LRRK2-G2019S Parkinson’s disease.

**Results:** The amount of PAK6 and 14-3-3γ is significantly different in patients with Parkinson’s disease compared to healthy subjects. Moreover, the amount of PAK6 also varies with the presence of the G2019S mutation in the LRRK2 gene. Although the generalized linear models show a low association between the presence of PD and PAK6, the kinase can be added in a broader panel of biomarkers for the diagnosis of Parkinson’s disease.

**Conclusions:** Changes of PAK6 and 14-3-3γ amount in plasma represent a shared readout for patients affected by sporadic and LRRK2-linked Parkinson’s disease. Overall, they can contribute to the establishment of an extended panel of biomarkers for the diagnosis of Parkinson’s disease.

## 3. Introduction

Parkinson’s disease (PD) is the second most common neurodegenerative disorder, affecting about 1-2% of people aged over 65 (1). Diagnosis of PD mainly relies on the clinical examination of patients’ motor symptoms, which typically start to appear when a considerable portion (about 60%) of dopaminergic neurons in the *substantia nigra pars compacta* is lost and nigrostriatal connectivity is irreversibly damaged (2). This late diagnosis and the consequent delay in the pharmacological treatment may in part explain why neuro-restorative/protective therapies have been failing so far in PD. Therefore, in the next future it will be crucial to find novel and specific biomarkers able to identify PD patients in their early clinical stages.

Remarkable technological advances have allowed the identification of several genes involved in PD (3). Among these, the one encoding leucine-rich repeat kinase 2 (LRRK2) plays a prominent role in the pathogenesis of both familial and sporadic PD (4-9). Up to date, over 100 variants have been ascribed to the *LRRK2* gene, of which only a few are causative of PD (6, 10, 11). In particular, the G2019S mutation is the most frequent, accounting for about 5% of familial cases and 1% of sporadic PD, although these percentages may vary according to the ethnic background (12). We have previously identified p21-activated kinase 6 (PAK6) as a strong interactor for LRRK2 (13, 14). PAK6 belongs to the group II PAKs (PAK4-6), which differ from group I PAKs (PAK1-3) for their sequence, mechanisms of activation and functions (15, 16). Collectively, PAK members play a major role in signal transduction and control a variety of cellular activities, including the dynamic cycle of polymerization and depolymerization of actin filaments as well as the rearrangement of microtubule networks (17-19). This aspect is of relevance in the central nervous system, where cytoskeleton remodelling is necessary for the development and refinement of dendritic spines and neuronal arborisation (20-22). Notably, also LRRK2 has been implicated in the regulation of actin dynamics, thus suggesting that PAK6 may work as an effector of LRRK2 (23-25). We have shown that PAK6 activity favours neurite outgrowth and the presence of LRRK2 is necessary to allow PAK6-mediated neuronal complexity (13). Both LRRK2 and PAK6 interact with 14-3-3s, a conserved family of dimeric proteins able to recognize and bind specific phospho-sequences on client partners, finally affecting their stability, functionality and subcellular localization (26-29). The phosphorylation of a cluster of serine residues located at the N terminal of LRRK2 and of Ser1444, is crucial to favour the binding of 14-3-3s to LRRK2 (30, 31). Likewise, also 14-3-3s can be phosphorylated at some critical Ser/Thr residues (Ser 58/59, Ser 184/186 and Ser/Thr 232, according to the isoforms) by different kinases (32-34). We showed that active/autophosphorylated PAK6 efficiently binds and phosphorylates 14-3-3γ at Ser59. Consequently, 14-3-3γ dimers dissociate from LRRK2 (35). We also showed that PAK6-mediated phosphorylation of 14-3-3γ can rescue neurite shortening in primary neurons derived from mice overexpressing the LRRK2 G2019S mutation, thus suggesting that the PAK6/14-3-3γ pathway may be part of the pathophysiology of LRRK2-related PD (35).

Here we evaluated whether plasma levels of PAK6 and 14-3-3γ can be exploited as candidate biomarkers for the detection of both sporadic and LRRK2-G2019S-linked PD. We show that both PAK6 and 14-3-3γ are differentially expressed in the brain and plasma of patients affected by PD compared to healthy subjects. Moreover, we found that PAK6 amount is also associated to the presence of the G2019S mutation, thus providing novel insights to unravel the role of PAK6 in PD.

## 4. Methods

### 4.1 Study participants

#### 4.1.1 Tissue samples

Post-mortem human caudate and putamen were obtained from Queen Square Brain Bank (London, UK). Tissues were collected under human tissue authority licence n° 12198. Limited sample demographics are listed in **Table1** and detailed in (36). A total of 14 samples were used, divided as follows: 4 LRRK2 G2019S-PD patients (LRRK2+PD+), 5 sporadic PD patients (LRRK2-PD+), and 5 age-matched controls (LRRK2-PD-). Samples were lysed in RIPA buffer containing 1% protease inhibitor cocktail (Sigma-Aldrich) and maintained at -80°C until use.

**Table 1:**
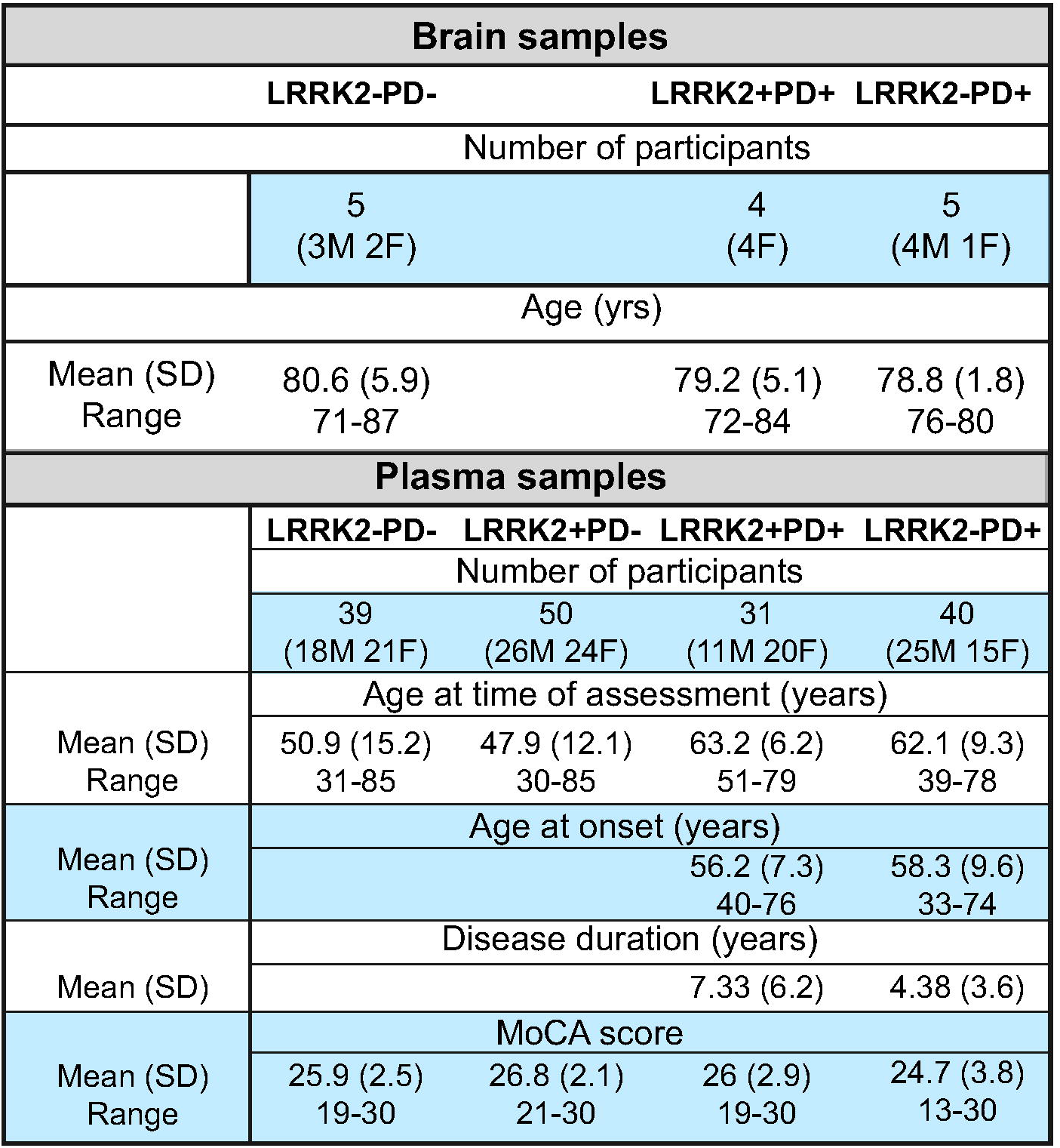
Demographic characterization of human brain and plasma samples. The table summarises the average number, sex and age of the brain samples’ donors employed in this study. The table also reports the MoCA score of all the donors of plasma samples used in this study as well as the mean age at disease onset and the mean disease duration for patients affected by PD.

#### 4.1.2 Plasma samples

Frozen plasma samples used in the analyses presented in this article were obtained from the LRRK2 Cohort Consortium (LCC) sponsored by the Michael J. Fox Foundation (MJFF). For up-to-date information on the study, visit www.michaeljfox.org./lcc. Case report forms and standard operating procedures can be found at https://www.michaeljfox.org/news/lrrk2-cohort-consortium. A total of 160 samples were used, divided as follows: 39 healthy subjects (LRRK2-PD-), 50 LRRK2 G2019S non-manifesting carriers (LRRK2+PD-), 31 LRRK2 G2019S-PD (LRRK2+PD+) and 40 sporadic PD (LRRK2-PD+). Samples were identified by a unique code and assigned to the 4 different groups after final measurements had been provided to the MJFF-LCC. Ethical review and approval was not required for the de-identified sample analysis in accordance with the local legislation and institutional requirements. The patients/participants provided their written informed consent to participate in this study.

### 4.2 Western blot

Protein concentration was measured using the Pierce™ BCA Protein Assay Kit following the manufacturer’s instructions (Thermo Scientific). 25µg of protein samples were resolved by electrophoresis on pre-cast 4–20% tris–glycine polyacrylamide gels (Bio Rad) and transferred to polyvinylidene difluoride membranes using a semi-dry Bio-Rad transfer machine (Trans-Blot™ Turbo TM Transfer System) with 1X Transfer Buffer (Bio-Rad) at 25V for 20 min. Membranes were incubated in 5% skimmed milk in Tris-buffered saline plus 0.1% Tween (TBS-T) for 1h at room temperature (RT), and then incubated overnight with primary antibodies diluted in 5% skimmed milk in TBS-T. The following primary antibodies were used: rabbit anti-PAK6 1:5000 (Abcam ab154752), rabbit anti-14-3-3γ 1:2000 (Thermo fisher PA5-29690) and mouse anti-GAPDH (CSB-MA000195, Cusabio, 1:3000). Membranes were subsequently rinsed and incubated for 1h at RT with the appropriate HorseRadish-Peroxidase (HRP)-conjugated secondary antibodies (Invitrogen). The visualisation of the signal was conducted using Immobilon™ Forte Western HRP Substrate (Millipore) and the VWR™ Imager Chemi Premium. Images were acquired and processed with ImageJ software to quantify the total intensity of each single band.

### 4.3 ELISA tests

The commercially available ELISA kits for human p21 protein (Cdc42/Rac)-activated kinase 6 (PAK6) (Mybiosource, MBS9318071) and for Human YWHAG (14-3-3γ) (Cusabio, CSB-EL026288HU) were used according to the manufacturing instructions. For each test, 50µl of plasma has been employed and the final concentration determined by adjusting for the used dilution. The detection range for the PAK6 ELISA kit extended from 0.25 to 8ng/ml, while for the 14-3-3γ ELISA kit it extended from 0.625 ng/mL to 40 ng/mL. Elisa kits were read and analysed with a Tecan Infinite 50 plate reader equipped with Magellan software 7.0.

### 4.4 Statistical analyses

The presence of outliers was determined by applying boxplot and histogram plots followed by Rosner’s test or generalized ESD many-outliers test (GESD) (37). Confirmed outliers were excluded from subsequent analyses. Analyses of demographic and clinical variables were performed with R studio (38). Given the statistical distribution of the data, investigated by the Shapiro-Wilk test, data were compared among the different groups by applying the non-parametric Kruskal-Wallis test followed by the Mann-Whitney post hoc test with Holm’s continuity correction (39). The statistical significance level was set to 0.05.

A Spearman’s correlation test was performed to study the presence of associations between PAK6 and 14-3-3γ in different subgroups.

Generalized linear models (GLMs) were implemented to assess a relationship between chosen independent and dependent variables (as described in the text). According to the need, the fit of the linear model was evaluated using the following indices (40) (41) i) McFadden’s index of explained variance (pseudo-R^2^) (42); ii) the study of the distributive normality of the model residuals (P_res_); iii) the Scaled Brier Score (sBS), which is a measure of overall accuracy and calculates the average prediction error (43); iv) Construction of the ROC (Receiver Operating Characteristic) curve and evaluation of the Area Under the Curve (AUC) and v) the Hosmer-Lemeshow test for fit between expected and estimated frequencies (χ ^2^_HL_ ; P _HL_) (44). The linear model was considered to fit the original data if the indices met the following criteria: i) the more pseudo-R^2^ is next to 1, the more the model is satisfactory; ii) normal distribution of residuals; iii) Brier score for a model can range from 0 (0%) for a perfect model to 1 (100%) for a non-informative model; iv) an AUC values >0.70 representing a moderately accurate model; v) a significant value indicating a bad model fit.

## 5. Data sharing

The data that support the findings of this study are available from the corresponding author upon reasonable request.

## 6. Results

### 6.1 PAK6 and 14-3-3γ expression in human brain tissues

We first compared the expression of PAK6 and 14-3-3γ in brain tissue obtained from LRRK2 G2019S-PD patients (LRRK2+PD+) (n=4), patients with sporadic PD (LRRK2-PD+) (n=5) and age-matched healthy subjects (LRRK2-PD-) (n=5). A demographic characterization of the population is reported in **Table 1**. We examined both sporadic and LRRK2 G2019S-PD patients to understand whether the expression of PAK6 and 14-3-3γ could offer a common readout for the two etiologically different pathologies. Indeed, while we have previously proved a functional interaction between LRRK2 and PAK6 (13, 35), a possible role of PAK6 in the onset of sporadic PD is currently unknown. As shown in **Fig.1A**, and quantified in the graphs, brain tissue derived from LRRK2+PD+ patients seem to present higher expression of PAK6 (p=0.127 **Fig. 1B**) and lower expression of 14-3-3γ (p=0.19 **Fig.1C**) compared to healthy subjects. Instead, LRRK2-PD+ patients tend to present lower expression of PAK6 and higher expression of 14-3-3γ (**Fig.1B-C**) compared to LRRK2-PD-subjects, (p=0.127 and p=0.19, respectively). However, in all these cases the difference does not reach statistical significance. Finally, LRRK2+PD+ patients present more PAK6 (p=0.048) and less 14-3-3γ (p=0.19) than LRRK2-PD+ patients. We also decided to consider the ratio between 14-3-3γ and PAK6, since it may provide higher specificity than single biomarkers. The 14-3-3γ/PAK6 ratio is significantly higher in LRRK2-PD+ patients compared to both LRRK2+PD+ patients and LRRK2-PD-subjects (p=0.048 for both comparisons) (**Fig.1D**).

**Figure 1:**
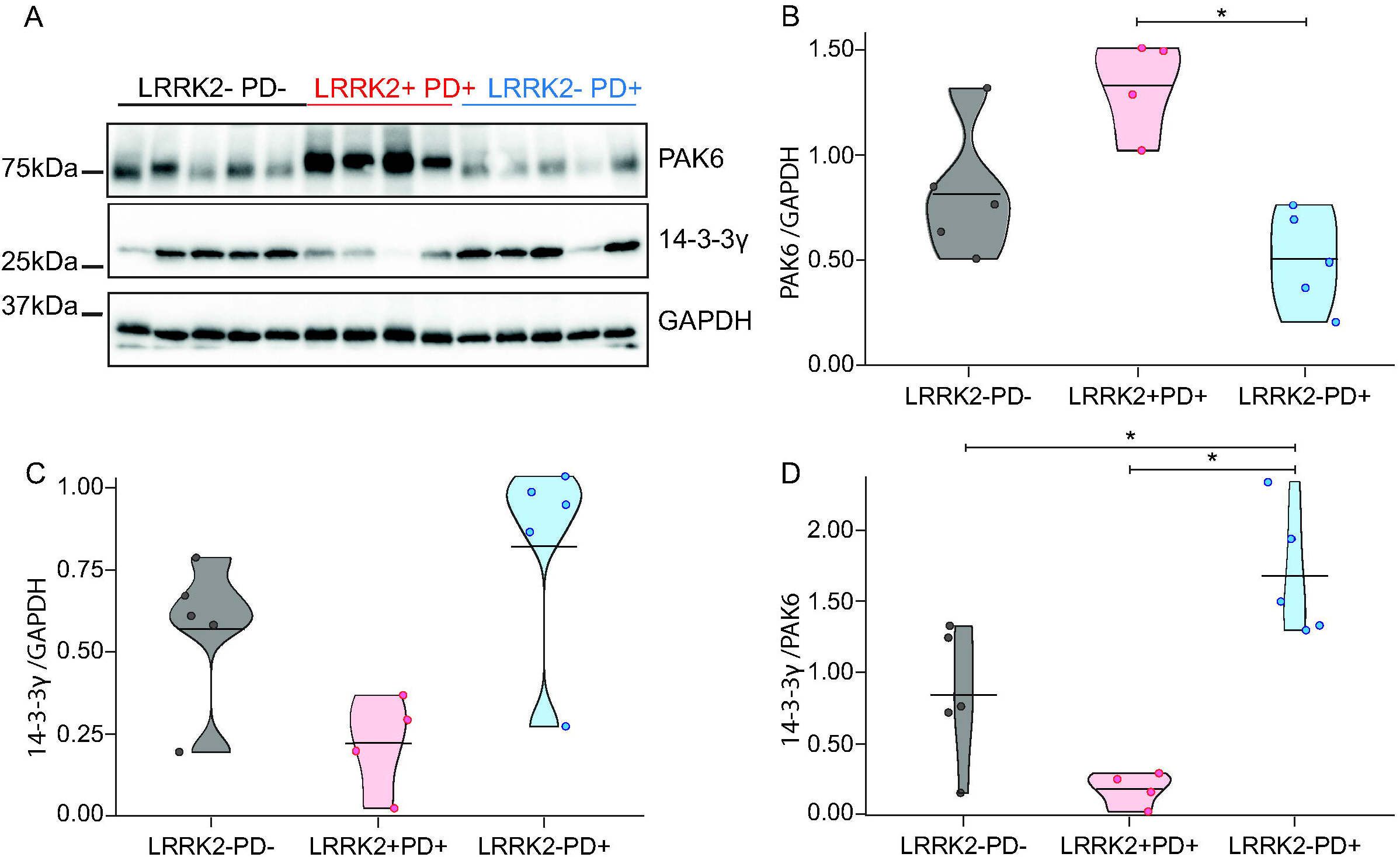
Expression of PAK6 and 14-3-3γ in human brains. (**A**) Western blot analysis of human LRRK2 G2019S and healthy controls brain tissues. (**B-D**) Relative quantification of band intensity of data shown in (**A**) normalised to GAPDH. Nonparametric Kruskal-Wallis test and pairwise comparisons using Mann-Whitney test with continuity correction were applied. *p values<0.05

These findings indicate that there is a differential expression of both PAK6 and 14-3-3γ in the brains of patients affected by sporadic and LRRK2 G2019S-PD.

### 6.2 Quantification of PAK6 and 14-3-3γ in human plasma

To test whether these differences are detectable also in tissues that are accessible from patients, we used a cohort of plasma samples. This allowed us to extend the number of samples and to include LRRK2 G2019S non-manifesting carriers, useful to distinguish if the amount of PAK6 and 14-3-3γ may correlate to the presence of the pathology or to the presence of the G2019S mutation in LRRK2.

We first delineated the demographic characteristics of our cohort, composed of healthy subjects (LRRK2-PD-), LRRK2 G2019S non-manifesting carriers (LRRK2+PD-), LRRK2 G2019S-PD patients (LRRK2+PD+) and sporadic PD patients (LRRK2-PD+). We noticed a significant difference among the groups as regards the age at the time of assessment. In particular, both LRRK2-PD- and LRRK2+PD-subjects are significantly younger than LRRK2-PD+ and LRRK2+PD+ patients (p<0.001). Instead, there is no significant difference between LRRK2-PD+ and LRRK2+PD+ patients nor for the age at disease onset (p=0.129), nor for disease duration (p=0.069). Finally, as for the MoCA score, a significant difference is shown between LRRK2-PD+ and LRRK2+PD-(p=0.017) (**Table 1**).

We next quantified the amount of PAK6, 14-3-3γ and the 14-3-3γ/PAK6 ratio in plasma samples by means of ELISA tests. We could not find any correlation of either PAK6, 14-3-3γ or the 14-3-3γ/PAK6 ratio with any of the demographic parameters analysed, with the only exception being for a correlation between the 14-3-3γ/PAK6 ratio and the age of assessment (**Table 2**).

**Table 2:** Correlation coefficient between PAK6, 14-3-3γ and 14-3-3γ/PAK6 and demographic parameters. The top part of the table summarises the correlation coefficient between PAK6, 14-3-3γ or the 14-3-3γ/PAK6 and the demographic parameters analysed in different groups. The bottom part of the table summarises the correlation test applied to verify the presence of any association between PAK6 and 14-3-3γ in different groups. **P-value <0.01.

| Correlation coefficient between PAK6, 14-3-3γ and 14-3-3γ/PAK6 and demographic parameters |  |  |  |  |
| --- | --- | --- | --- | --- |
| All subjects | Age at assessment | Age at onset | Disease duration | MoCA score |
| PAK6 | -0.17 | -0.16 | 0.01 | 0.10 |
| 14-3-3γ | 0.23 | 0.06 | 0.20 | -0.08 |
| 14-3-3γ/PAK6 | 0.31** | 0.09 | 0.16 | -0.12 |
| **LRRK2-PD-** |  |  |  |  |
| PAK6 | -0.08 |  |  | -0.24 |
| 14-3-3γ | 0.26 |  |  | 0.04 |
| 14-3-3γ/PAK6 | 0.32 |  |  | 0.04 |
| **LRRK2+PD-** |  |  |  |  |
| PAK6 | -0.03 |  |  | 0.16 |
| 14-3-3γ | 0.10 |  |  | -0.16 |
| 14-3-3γ/PAK6 | 0.18 |  |  | -0.14 |
| **LRRK2+PD+** |  |  |  |  |
| PAK6 | -0.01 | -0.03 | 0.05 | -0.04 |
| 14-3-3γ | -0.10 | -0.31 | 0.17 | 0.13 |
| 14-3-3γ/PAK6 | -0.06 | -0.25 | 0.14 | 0.15 |
| **LRRK2-PD+** |  |  |  |  |
| PAK6 | -0.27 | -0.16 | -0.13 | 0.12 |
| 14-3-3γ | 0.24 | 0.25 | 0.20 | -0.09 |
| 14-3-3γ/PAK6 | 0.30 | 0.28 | 0.21 | -0.18 |
| Correlation coefficient between PAK6 and 14-3-3γ in plasma samples |  |  |  |  |
| LRRK2-PD- | 0.21 |  |  |  |
| LRRK2+PD- | -0.07 |  |  |  |
| LRRK2+PD+ | 0.15 |  |  |  |
| LRRK2-PD+ | -0.17 |  |  |  |
| LRRK2- | -0.05 |  |  |  |
| LRRK2+ | 0.02 |  |  |  |
| PD- | 0.02 |  |  |  |
| PD+ | 0.08 |  |  |  |

As shown in **Fig.2**, there is a significant difference only between LRRK2-PD+ and LRRK2+PD-subjects for PAK6 (p=0.001) and for the 14-3-3γ/PAK6 ratio (p=0.001) (**Fig.2A and 2C**), while there is no difference among the 4 groups as for the values of 14-3-3γ (**Fig.2B**). However, we noticed that PD patients (both LRRK2-PD+ and LRRK2+ PD+) seemed to have less PAK6 and more 14-3-3γ compared to their respective controls (LRRK2-PD- and LRRK2+PD-), similar to what we had observed in the brain of sporadic PD patients (**Fig.1**). We also noticed that subjects with the G2019S mutation in LRRK2 gene tended to have more PAK6 compared to subjects without the mutation, once more reflecting our previous observations in the brain.

**Figure 2:**
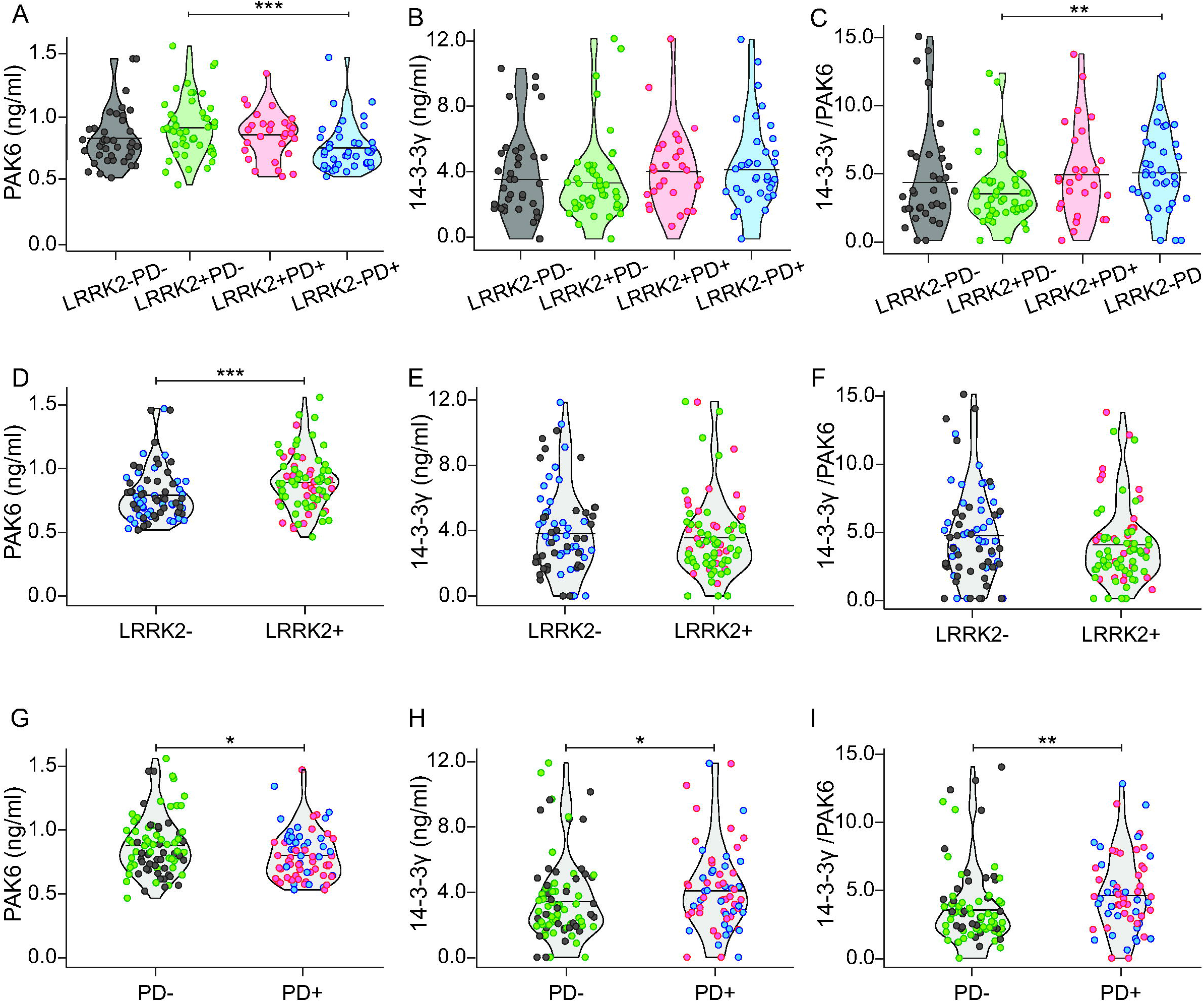
Distribution of PAK6, 14-3-3γ and 14-3-3γ/PAK6 among the groups. Graphical distribution of the amount of PAK6 (**A**), 14-3-3γ (**B**) and 14-3-3γ/PAK6 (**C**) in plasma samples of the 4 groups. Comparison of PAK6 (**D**), 14-3-3γ (**E**) and 14-3-3γ/PAK6 (**F**) amount in plasma samples of LRRK2-subjects and LRRK2 subjects. Comparison of PAK6 (**G**), 14-3-3γ (**H**) and 14-3-3γ/PAK6 (**I**) amount in plasma samples of PD-subjects and PD+ patients. Nonparametric Kruskal-Wallis test and pairwise comparisons using Mann-Whitney test with continuity correction were applied. *p values<0.05 **p values<0.01, ***p values <0.001

Altogether, these data suggest that the amount of PAK6 (and to a lesser extent 14-3-3γ) is influenced by the presence of the mutation and/or the presence of the disease.

### 6.3 PAK6 amount changes with disease condition and genetic background

We then stratified our data by grouping the subjects according to their genetic background; namely LRRK2-PD- and LRRK2-PD+ (indicated as LRRK2-) and LRRK2+PD-with LRRK2+PD+ (indicated as LRRK2+). As shown in **Fig.2D-F**, PAK6 is significantly higher in subjects with the mutation compared to subjects without the mutation (p<0.001). Instead, there is no significant difference either for 14-3-3γ (p=0.484) or for the 14-3-3γ/PAK6 ratio (p=0.156) between the two groups.

Likewise, we grouped together data from subjects according to their disease condition; namely LRRK2-PD- and LRRK2+PD-(indicated as PD-) and LRRK2-PD+ with LRRK2+PD+ (indicated as PD+). As shown in **Fig.2G-I**, PAK6 is significantly higher in PD-subjects compared to PD+ patients (p=0.023). On the opposite, 14-3-3γ and the 14-3-3γ/PAK6 ratio are higher in PD patients compared to PD-subjects (p=0.036 and p=0.004, respectively). We therefore wondered if the two proteins in the plasma were inversely correlated. However, no correlation has been found in any of the groups analysed (**Table 2**).

All together these observations indicate that there is a relation between PAK6 (and to a lesser extent 14-3-3γ) and the probability of having the disease or the mutation.

### 6.4 Generalized linear model for PD

We therefore applied generalized linear models to investigate the relationship between the probability of having PD (dependent variable) and a group of variables selected as predictors (namely: PAK6 amount, 14-3-3γ amount, 14-3-3γ/PAK6 ratio, age at time of assessment, gender and MoCA score). According to our data, PAK6 is the only variable able to influence the probability of having PD (β=-1.88; P=0.02) (**Fig.3A**). The estimated model appears to be very informative (sBs=0.04) and the fit between the predicted and estimated frequencies is also satisfactory (χ2= 4.29; P=0.829). However, the receiver operating characteristic (ROC) curve, which allows the evaluation of the predictive ability, shows that the estimate of the area under the curve (AUC) is 0.61, thus indicating poor accuracy prediction (61%).

**Figure 3:**
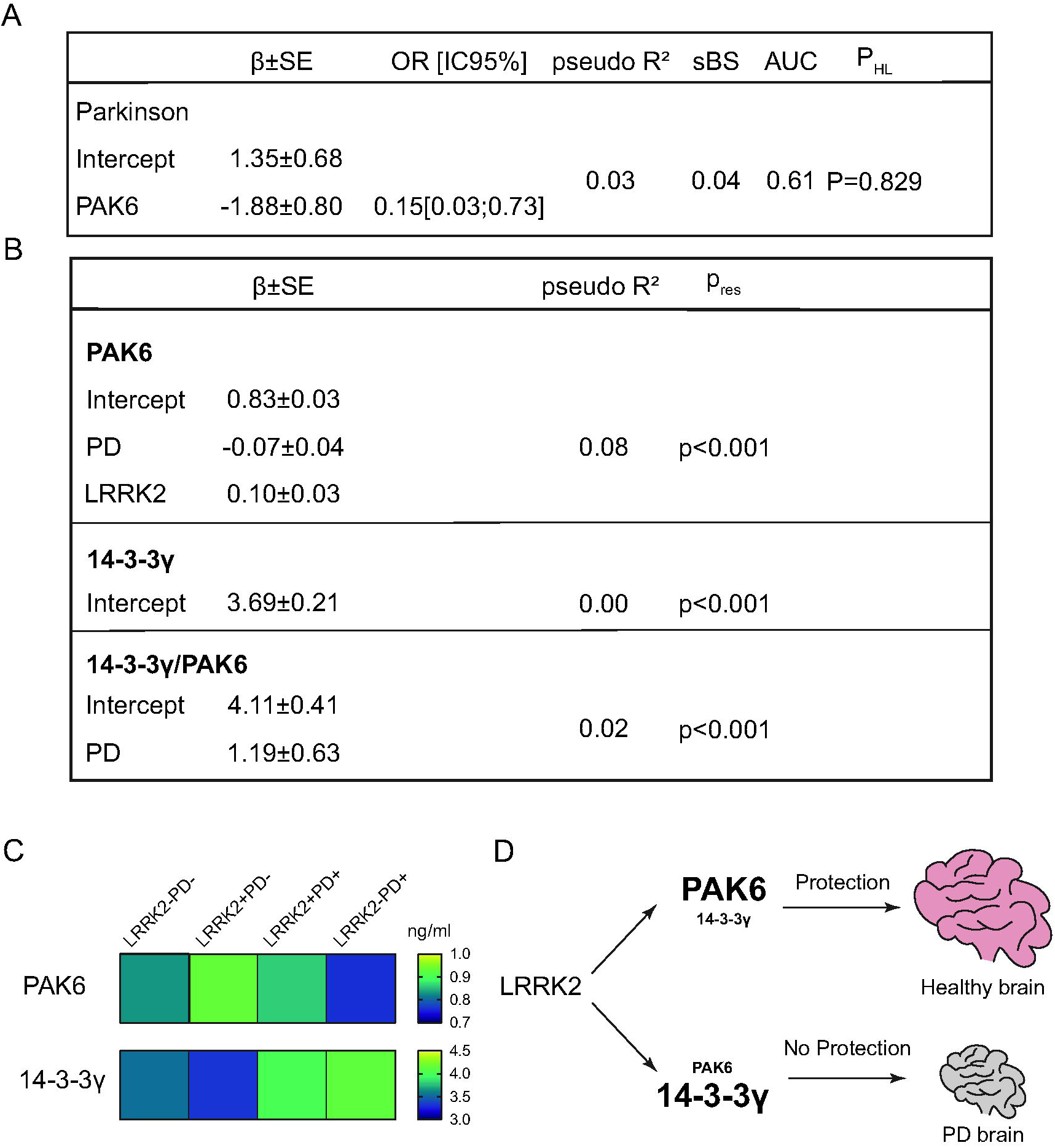
Generalized linear models for the presence of PD, PAK6, 14-3-3γ and 14-3-3γ/PAK6. Generalized linear models, showing the relationship between the PAK6 amount and the probability of having Parkinson’s disease (**A**) and the relationship between PAK6, 14-3-3γ and 14-3-3γ/PAK6 and different parameters as described in the text (**B**). Outcomes are displayed as estimate of regression coefficient with Standard Error (β ± SE); McFadden’s index of explained variance (pseudo-R2); Scaled Brier Score (sBS); Area Under the Curve (AUC); p-value of the Hosmer-Lemeshow test (PHL) or p-value of the Shapiro Wilk test of model residuals (P_res_). Significance was established at P<0.05*.(**C**) Gradient colour representation of PAK6 and 14-3-3γ amount in plasma samples. (**D**) Schematic underlying the protective role of PAK6.

We also applied a similar linear model analysis to evaluate which of the predictors taken into consideration (PAK6, 14-3-3γ, 14-3-3γ/PAK6, age at time of assessment, gender, presence/absence of LRRK2 mutation, presence/absence of disease, age at disease onset, disease duration, MoCA score) influence the amount of PAK6, 14-3-3γ or the 14-3-3γ/PAK6 ratio. While none of the variables seem to influence the amount of 14-3-3γ, the presence of the disease shows a positive relation with the 14-3-3γ/PAK6 ratio (β = 1.19; P = 0.05) and a negative relation with the amount of PAK6 (β = - 0.07; P = 0.041). Moreover, there is a positive relation between the amount of PAK6 and the presence of the mutation (β = 0.10; P = 0.005). However, the model does not fit the data well, with a very low McFadden’s index of explained variance (pseudo-R^2^=0.08) and a non-normal distribution of model residuals (P_res_<0.001) (**Fig.3B**).

All together these statistical analyses confirm our experimental observations, according to which the amount of PAK6 (and to a lesser extent the 14-3-3γ/PAK6 ratio) in plasma is related to both the presence of the G2019S mutation in the LRRK2 gene and to the presence of the disease.Although the predictive power is not high, we can conclude that PAK6 and 14-3-3γ can be used in concert with other biomarkers to define a novel panel of biomarkers for the diagnosis of PD.

## 7. Discussion

In the present study, we explored PAK6 and 14-3-3γ as novel biomarkers for the diagnosis of PD. We have previously shown that PAK6 is functionally related to LRRK2, and that it is aberrantly activated in the brain of PD patients (13), thus supporting the idea that PAK6 may play a role in the onset of PD. However, the relation between PAK6 and the aetiology of sporadic PD still needs to be investigated. Likewise, several observations have been linking different isoforms of 14-3-3 proteins to PD. First, 14-3-3s are present in Lewy bodies and are known to interact and co-localize with several proteins involved in PD pathogenesis, including LRRK2, α-Syn, and Parkin (28, 45). Secondly, 14-3-3 isoforms have been shown to exert a protective role in models of PD, both in vitro and in vivo (46-48); finally, aberrantly phosphorylated 14-3-3s have been found in brain lysates from PD patients (49).

Here, we first evaluated the expression of PAK6, 14-3-3γ and 14-3-3γ/PAK6 ratio in brains of patients affected by sporadic PD, LRRK2 G2019S-PD and healthy subjects. We reported that there was a differential expression of both PAK6 and 14-3-3γ in the two groups of PD patients.

We therefore extended our analyses to a larger cohort of plasma samples and included non-manifesting carriers, whose presence is important to understand whether changes of PAK6 and/or 14-3-3γ amount could be related to the presence of the disease or to the mutation itself. To the best of our knowledge, PAK6 has not been detected in biological fluids, while the detection of 14-3-3s has been performed mainly on samples of cerebrospinal fluid, although some attempts have been made to detect 14-3-3s in blood-derived biosamples (50-54). In general, the trend of both PAK6 and 14-3-3γ in plasma samples was in line with the one observed in brain samples, thus suggesting that plasma could represent an appropriate biofluid to investigate the relevance of these two proteins in the context of PD. We observed that there was a significant difference in the amount of PAK6 and 14-3-3γ/PAK6 ratio between LRRK2-PD+ and LRRK2+PD-subjects, while there was no significant difference between the 4 groups for 14-3-3γ. This prompted us to consider that PAK6 had a stronger significance compared to 14-3-3γ. In particular, the fact that it showed a significant difference between LRRK2-PD+ and LRRK2+PD-patients, which differ both for the genetic background and the presence of the disease, allowed us to speculate that PAK6 could depend on these two variables. We therefore decided to group our samples based on their genetic background or on their disease condition. According to these analyses, we observed that the amount of PAK6 and 14-3-3γ were respectively lower and higher in patients affected by PD compared to healthy subjects; moreover, PAK6 amount was higher in subjects with the mutation in the LRRK2 gene compared to subjects without the mutation (**Fig. 3C**). These results have been further confirmed with the application of GLMs, which validated the reciprocal inverse relation between PAK6 and the probability of having the disease, as well as the positive connection between PAK6 and the mutation in LRRK2. However, the poor predictive value estimated by the ROC curve (0.61) prevents us from concluding that PAK6 values in the plasma can distinguish people affected by PD from healthy subjects. Finally, we could not find any significant correlation between PAK6 or 14-3-3γ with any of the demographical parameters analysed. In general, we observed that PAK6 shows a stronger relation with PD compared to 14-3-3γ. Indeed, 14-3-3γ works as a downstream target in the PAK6-LRRK2 axis, therefore the pathological effects may be somehow diluted. Moreover, 14-3-3 proteins are involved in a myriad of cellular processes, so their role in PD may be less specific than PAK6. Nevertheless, the 14-3-3γ/PAK6 ratio can still provide a useful indication to discriminate PD patients from healthy controls, especially if used in concert with other candidate biomarkers for PD (55, 56). In this work we evaluated the -γ isoform because it interacts with PAK6 with higher affinity compared to other isoforms (35). However, we cannot exclude that other 14-3-3s, which have been shown to exert a protective role in PD (29, 48, 57), can work as potential biomarkers in PD.

The relation between PAK6 and the mutation in LRRK2 strengthens our previous studies and poses new questions. Most LRRK2 mutations (including the G2019S) induce an increase of its kinase activity, which finally ends up in a cytotoxic effect (58, 59). Indeed, the measurement of LRRK2 activity has been proposed as a possible biomarker readout in several studies (60-65). We have previously shown that LRRK2 is crucial for PAK6-mediated neuronal complexity and that LRRK2 alone is not sufficient to induce PAK6 activation *in vitro*. Moreover, we proved that overactivated PAK6 can rescue G2019S-LRRK2-induced neurite shortening via phosphorylation of 14-3-3γ. This protective mechanism is likely mediated by the ability of PAK6 kinase activity to lower LRRK2-substrates hyperphosphorylation (14). In line with this model, PAK6 is hyperactivated in brains of both G2019S-LRRK2 PD and sporadic PD patients (13, 35). Here we show that PD patients present less PAK6 than healthy subjects, and that subjects with the G2019S LRRK2 mutation have an increased amount of PAK6 compared to subjects without the mutation. These observations allow us to speculate that the presence of the G2019S mutation in LRRK2 induces the expression of PAK6 as a self-protecting attempt to buffer the activity of the mutated kinase, likely through an increased phosphorylation of 14-3-3γ. We further hypothesize that if the G2019S LRRK2-induced endogenous overexpression of PAK6 is not sufficient, LRRK2 hyperactivation would not be counteracted and the subject may eventually develop PD (**Fig. 3D**). However, while this provides a possible explanation for patients affected by the G2019S mutation in LRRK2, it does not explain why patients affected by sporadic PD still show a tendency to present less PAK6 in their plasma compared to healthy subjects. In light of these data, we propose PAK6 as a convergent player in the pathogenesis of both sporadic and G2019S-LRKK2 PD, an intriguing hypothesis that will need further investigation.

In conclusion, we suggest that PAK6 may be likely added in a hypothetical future panel for the diagnosis of PD. However, further studies with different LRRK2 variants will help us to elucidate how mutations in LRRK2 induce an increase in the expression of PAK6 and how the overall expression of PAK6 is crucial to determine the clinical development of the disease. Moreover, if PAK6 should be confirmed to be at the crossroads between sporadic and G2019S-LRRK2 PD, this may open new perspectives in the search of novel therapeutic strategies.

## 8. Acknowledgment

University of Padova to support LC as assistant professor and the IRCCS San Camillo Hospital in Venice, Italy. The LRRK2 Cohort Consortium, which is coordinated and funded by The Michael J.Fox Foundation for Parkinson’s Research. The investigators within the LCC contributed to the design and implementation of the LCC and/or provided data and/or collected biospecimens, but did not necessarily participate in the analysis or writing of this report. The full list of LCC investigators can be found at www.michaeljfox.org/lccinvestigators. We also would like to thank Dr. Giorgio Arcara for his insightful comments on the manuscript.

## 9. Authors’ roles

EGi and LC designed the study, performed and optimised the experiments, interpreted data, and wrote the manuscript; LM performed statistical analyses; LIa performed western blot experiments; VG and LIo helped with the preparation of the samples. RBan provided human brain samples. AA, LB, RBar, NP and EGr have contributed to manuscript editing and optimization. All authors have read and approved the final manuscript.

